# DFT Modelling of the Electronic Structure and Stabilities of (Sb_2_O_5_.nH_2_O) Clusters towards Cysteine, Glucose and Trypanothione Complexation

**DOI:** 10.1101/2020.06.29.177360

**Authors:** Hassan Rabaa, Andriy Garfov, Dage Sundholm

**Author notes:** **Dedicated to Emeritus Prof. Jean-Yves Saillard from Rennes 1 University (France)**. Corresponding author **Email** (**MA**). **Supporting information is given via a link at the end of the document.**.

## Abstract

A large series of dipeptides containing sulfur groups and antimony Sb^V^ were modeled to understand their inhibitor activity against Leishmaniasis. The trypanothione reductase (TR), which acts as a reducing agent in several vital processes, is responsible for maintaining the parasite’s cellular thiol redox balance. The antimonic Sb^V^ acid (Sb_2_O_5_·nH_2_O) is being evaluated as a drug with inhibitory activity against Leishmaniasis. In the present work, we investigated the inhibitory effect of antimony oxide on (TR) activity modeled as a substrate by probing two model clusters in gas phase and continuum water medium: **A** [(Sb_2_O_10_H_8_)]^−2^ coordinated to cysteine, and **B** [Sb_7_O_28_H_21_] coordinated to trypanothione, including glucose adduct. We report here density functional theory (DFT and DFT-D3) using (B3LYP/LANL2DZ and (TPSS/def2TZVP) results on the binding energy of cysteine and trypanthione complexed to these clusters as possible sites promoting the inhibition process. Upon viewing the results of the computational studies of cluster models and theoretical thermochemistry data for receptor-substrate interactions, identification of ligand-cluster interactions helps to unravel the mechanism of inhibition. The acidity of (Sb_2_O_5_,nH_2_O) leads to great cluster-dipeptide passivation. The electrostatic forces between cluster interface and dipeptide interaction present relevant inhibition effects through proton transfer or mobility from the different amine and ketone groups. The reactivity differences come from the unoccupied lone pairs 5p_Sb_ which lie at higher energy but remain available to make a good interaction with the lowest orbital p nitrogen in NH_2_, and in the CO groups substrate fragment in the zone HOMO-LUMO. Further cluster stability comparisons show a lower Gibbs free energy in **B3** (**B**/trypanthione/glucose) (18 – 30 kcal/mol) at both used level in this study and gives good accurate intramolecular interactions, confirmed by the use of the dispersion-corrected density functional (DFT-D3). Given the dipeptide H-mobility and the (Sb_2_O_5_,nH_2_O) cluster acidity, (donor-acceptor duality), the system is predicted to be potent cluster of the inhibitors by endothermic and spontaneous reaction requiring 3.10 kcal/mol in aqueous medium.

## Introduction

For more than a century, Sb^V^ antimonial drugs, for chemotherapy of Leishmaniasis, have been routinely prescribed [1–9]. Little is known about their toxicity [2b] and pharmacological mechanisms, and yet they continue to be the first choice drugs for this disease. As the primary drug used in the treatment of Leishmaniasis, studies into its toxicity and pharmacological mechanisms have yet to be completed. In the search for agents with higher therapeutic indices, a number of other antimonial pentavalent compounds were discovered in the 1920s [6–13] and used as drugs to treat Leishmaniasis. Antimony is often considered as a semimetal, with most common oxidation states being trivalent Sb^III^ and pentavalent Sb^V^ [1]. In the literature, several dipeptidyl compounds based on a nitrile/sulfur scaffold were synthesized and probed for their inhibitory activity against Leishmaniasis. Recent research reported that trypanthione acts as a reducing agent in several vital processes and is also responsible for maintaining the parasite’s cellular thiol redox balance [4–7]. Different studies have suggested the importance of genes involved in trypanothione metabolism and drug transport [9–13] in conjunction with trypanothione as a possible treatment option with potential antileishmanial activity. Trypanothione T(SH)_2_ is synthesized from glutathione and spermidine by trypanothione synthetase (TS) and is further reduced by trypanothione reductase (TR) [9–13]. Fairlamb and co-workers have shown that pentavalent antimony, the main drugs used against Leishmaniasis, in vivo interfere with trypanothione metabolism by inducing rapid efflux of intracellular T(SH)_2_ and by inhibiting TR in intact cells [9].

Trypanothione Reductase (TR) is known as the key enzyme in Leishmaniasis infection [9–13]. Although this enzyme is thought to be a potential drug target, the development of an accurate TR inhibitor is still required [9]. Trypanothione participates also in crucial thiol–disulfide exchange reactions and serves as an electron donor in several metabolic pathways from synthesis of DNA precursors to oxidant detoxification inhibitory activity against Leishmaniasis reduced inside. The amastigote to Sb^III^, interferes *in vivo* with the T(SH)_2_ metabolism by inducing rapid efflux of intracellular T(SH)_2_ and inhibiting TR in intact cells [9–13]. Thus, antimony compounds are used as antibacterial agents [10]. Several metal complexes have been tested on a variety of trypanosomatids, and were shown to be active on different molecular targets [10–13]. Drug-Vectors Nanostructures (DVN) and encapsulated polymers studies emerge as well, as was reported by Grafov et al, who synthesized compatible (DVN) using Sb^V^ and glucantime, and published under the available patent (# BR 10 2015 2013 0236180) [16]. Goodwin et al, were the first to propose that Sb^V^ could act as a pro-drug that has to be converted into active Sb^III^ [17].

In this report, the model cluster was built from Sb_2_O_5_ oxide X-ray data [18]. Different theoretical studies were published in this area [19–21]. In this account, two cluster models of hydoxylated antimony oxide (Sb_2_O_5_,nH_2_O)/dipeptide have been computationally investigated at the density functional theory (DFT) levels of theory using Gaussian G16 [22] and Turbomol 7.3 [23] packages: **A** [(Sb_2_O_10_H_8_)]^−2^ coordinated to the cysteine/glucose, and **B [**Sb_7_O_28_H_21_] coordinated to trypanthione/glucose [20]. To elucidate all electronic factors influencing the dipeptide/(Sb_2_O_5_,nH_2_O) clusters stability during the inhibition cluster process, different quantum tools were employed for water simulating at the DFT level. LANL2DZ basis set was used for all atoms and 6-311G**(d.p) for main atoms instead Sb [24–27] and probed by TPSS/def2TZVP [28–31]. The (continuum solvation model) SMD model, was used [32–33]. Analysis of ΔG calculations, and basis set super position error (BSSE) [34] as well as the binding energies were probed. Analyses of the interaction of hydroxylated antimony oxides clusters with dipeptide in vacuum or wet medium, were determined. Theoretical parameters of dipole moment (μ), E_HOMO_, E_LUMO_, HOMO–LUMO gap analysis, binding energy (BE), molecular electrostatic potential (MEP) [35]. Thermodynamic properties and Mulliken atomic charges to seek out the essential molecular requirements for the inhibitory activity upon targeting cysteine or trypanothione. The electrochemical corrosion phenomenon in acidic medium occurs in the liquid phase, and it is well known that the inhibitory molecules in medium act otherwise than in a vacuum, this is why it is relevant to include the solvent effect in the computations. Dispersion-corrected density functional theory calculations (GD3BJ) using TPSS/Def2-TZVP as functional and basis sets were performed to explore the main intramolecular interactions present in this molecular modeling [36]. In the text, (DFT-D3) is used as an acronym for TPSS/Def2-TZVP (DFT-GD3BJ).

### Computational Methodology

In an effort to assess the generality of the attraction hypothesis cluster/dipeptide, the present study is focused on the electronic structures and the thermodynamic stability using quantum chemistry tools. The model cluster (Sb_2_O_5_,nH_2_O) were optimized at DFT/B3LYP as implemented in the Gaussian 16 (G16) [22] and Turbomol 7.3 packages [23]. LANL2DZ basis set was used for all atoms and 6-31g**(d.p) for main atoms instead Sb [24–25]. And also all models were optimized at DFT-D3 for all stable structure confirmation [25–31].

The cluster model [(Sb_2_O_10_H_8_)]^−2^ **A** was computed in gas phase and SMD/water [32–33] after probing various cluster/dipeptide conformation. The large neutral cluster [Sb_7_O_23_H_19_)] **B** is shown in **Fig 1**. Therefore, in term of simulating water environment in **B**, we add randomly different protons in terminal oxygens for keeping the total cluster neutrality. The Cartesian coordinates of the molecular structures optimized at B3LYP/LANL2DZ and TPSS/def2-TZVP levels of theory are given as Supporting Information.

**Fig 1.**
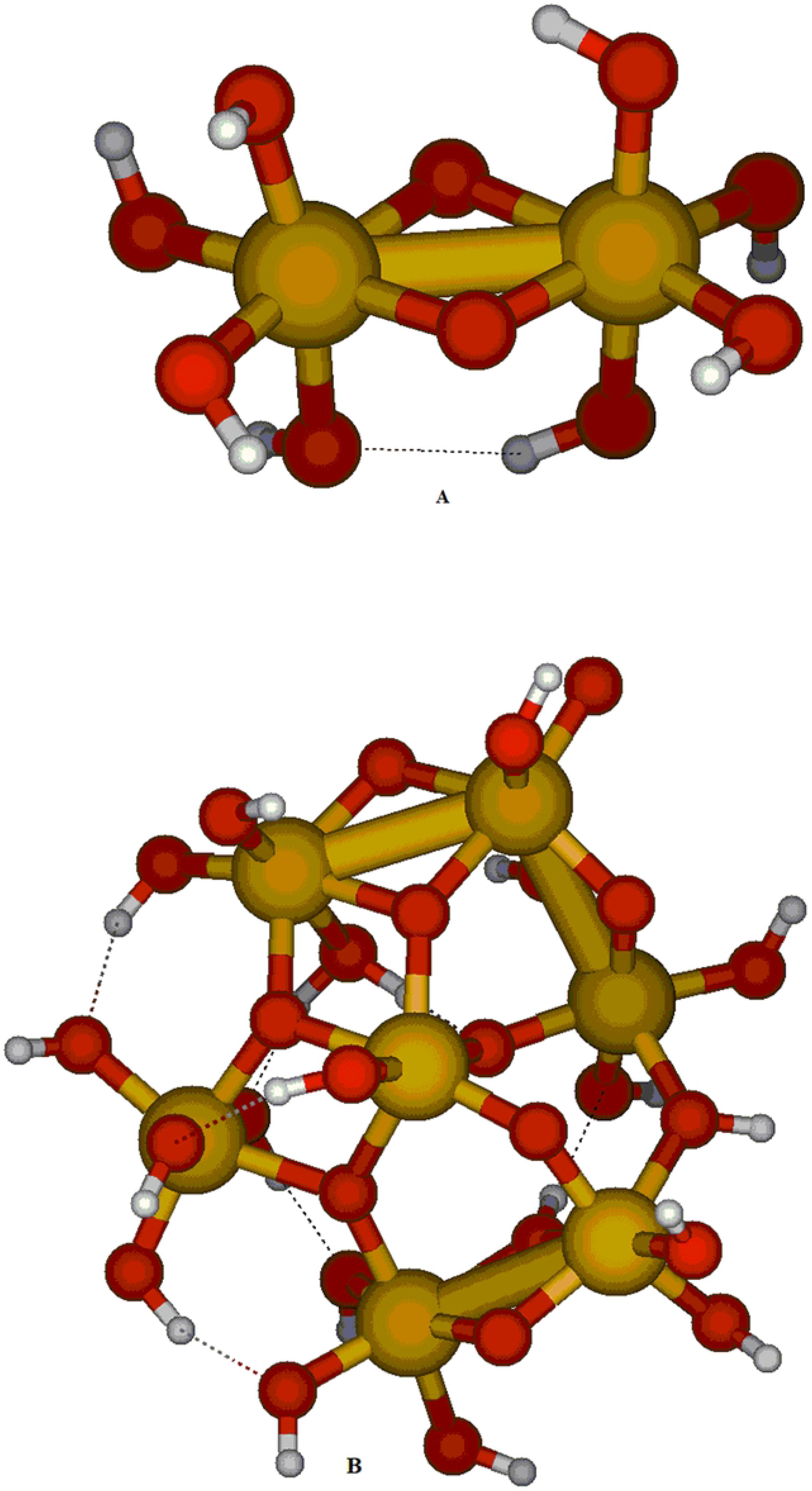
Optimized molecular structures of cluster [(Sb_2_O_10_H_8_)]^−2^ **A** (right) and [(Sb_7_O_28_H_21_)] **B** (left) at B3lyp/Lanl2dz. In **A,** Sb atoms are presented by purple or yellow ball; O: red ball and gray or white ball: Hand in **B,** Sb atom is shown in yellow, and the O atoms in red and H atoms in white.

All considered dipeptide/cluster structures in this study were optimized in the following way. Added molecules on the cluster were placed in coplanar planes in such a way that their main axes is parallel. Vertical separation distance of 4.1 angstroms was set for the energy optimization and in plane displacement was optimized. The optimized generated tautomers including glucose and cysteine, or trypanothione complexation, were converged with positive frequencies. Harmonic frequencies were calculated for each complex, to verify the stationary point minima on the potential energy surface. A singlet spin state was assumed and no symmetry constraints were imposed. The total energy calculation of certain clusters with or without basis set super position error (BSSE) were performed [34].

Electronic structures, Mulliken charges, HOMO-LUMO, dipolar moment, and solvent effects were discussed to support the Sb-O-H-ligand interactions in these cluster models. Dispersion-corrected density functional theory calculations (DFT-D3) were performed for the main intramolecular interactions present in this molecular modeling [36]. In addition, the molecular electrostatic potential (MEP) profile, considered as the crucial tool to grasp the molecular interactions was carried out. Cluster models have drawn great attention, due to their fundamental interest in basic research, as well as the possibility of constructing nanostructured materials using clusters as building blocks [37–39].

## RESULTS AND DISCUSSION

### Preliminary calculations

First optimized tautomer of [(Sb_2_O_10_H_8_)]^−2^ **A** converges in several conformers, (**a, b, c** and **d**) are shown in **SI-Fig. 1**. The Cartesian coordinates of the optimized structures are given as Supporting Information (**SI**). Metric parameters obtained from B3LYP/LANL2DZ calculations are presented in **Table 1**. Most optimized Sb-O bond lengths vary from 1.990 Å to 2.006 Å including pentavalent antimony Sb^V^ in aqueous systems. O-Sb-O bond angles ranging from 82° to 96°, deviate slightly from the experimental values [18]. In **Table 1**, the three optimized (**a, b** and **c**) tautomers present almost the same molecular structures as drawn in **SI-Fig. 1**, unlike in **d,** where it converges with an isolated OH_2_ molecule. The HOMO-LUMO gap energy ranging between 2.10 - 2.50 eV, is consistent with good stability. No substantial changes in bond length were seen in water simulating (see last row in **Table 1**). Calculations of the Gibbs free energy give **a** as the most stable structure, whereas **b**, **c** and **d** remain more higher by 14 kcal/mol. Embedding the complexes in SMD/water did not significantly change the relative Gibbs free energy. Structure a still is energetically the lowest one, and **d** the highest conformation by 18 kcal/mol. The **d** molecular structure is unfavorable due to the Sb=O bond formation and the protonated bridging oxygen, as shown by the relative Gibbs free energies given in **Table 1**. By comparison to normal DFT, small changes was seen for **a** computed at DFT-D3, unless Sb-Sb bonds were decreased from 3.143 Å to 3.041 Å (table 1, first column). Binding energy remains quite similar.

**Table 1.** Calculated bond averages for optimized tautomers of [(Sb_2_O_10_H_8_)]^−2^ (**a**, **b**, **c** and **d**) in gas phase at B3LYP/LANl2DZ and in parentheses for the most stable (**a**) at **TPSS/Def2-TZVP (DFT-GD3BJ**: (O_b_ (Bridge) and O_t_ (Terminal oxygens). Gibbs free energy in (kcal/mol) calculated in gas phase and in SMD=Water. Distances are in Angstrom. Angles are in degrees.

| Distance in Tautomer | **a** | **b** | **c** | **d** |
| --- | --- | --- | --- | --- |
| Sb<sub>1</sub>-Sb | 3.143 ( 3.041) | 3.298 | 3.622 | 3.958 |
| Sb-$\text{O}_t$ | 2.037 (2.011) | 1.908 | 1.987 | 1.888 |
| Sb-$\text{O}_b$ | 2.050 (1.995) | 2.144 | 2.111 | 1.952 |
| Sb-$\text{O}_H$ | 1.999 (1.895) | 2.020 | 1.991 | 1.997 |
| O-H | 0.976 (0.918) | 0.970 | 2.016 | 2.016 |
| $\text{O}_b$-Sb-$\text{O}_b$ | 94.4 (98) | 95.6 | 82.2 | 96.1 |
| $\Delta G$ (kcal/mol)/Gas Phase | 0. (12) | 12.1 | 14.50 | 13.80 |
| $\Delta G$ (kcal/mol)/SMD=Water | 0. (09) | 11.48 | 12.78 | 18.73 |

Looking closer at the molecular structure of **a,** every antimony atom shows six bonds, as expected in a classical ML_6_ octahedron with M= Sb and L= O). Different bonding types of oxygen were observed: bridged metal-metal in **a**, mono-bridged metal-metal in **b**, and in **c** and **d** bridged protonated oxygen. Most Sb-O bonds are in the range of 1.999 to 2.061 Å. In addition, we do not observe any modification of O-Sb-O angles on inclusion of solvent. Angles are ranging from 76.5 to 94.4° coorobore the X-ray structure [18]. Moreover, in gas phase, the Gibbs free energies range from 12.10 - 13.80 kcal/mol yielding **a** as the most stable conformer with bridged metal-metal and metal-oxygen bonds as both od levels of theory

The second attempt of geometry optimization was of the glucose complexation of (**a**, **b**, **c** and **d**) **(SI-Fig 2)** yielding **e**, **f**, **h** and **g** complexes as shown in supplementary information. We focus particularly in their binding energies upon cyclic glucose addition. Most relevant optimized structures in gas phase of (**e, f, h** and **g**), calculated at B3LYP/LANL2DZ, are shown in **SI**-**Table 1**. The Cartesian coordinates of the optimized structures are given as **SI**. As results, we observe extra inter-connectivity of non-bonding OH between cluster-cysteine). These O-H bonds pointing towards the cluster and making weak interaction type van der Waals interactions, range from 1.581 to 1.872 Å. The binding energy is decreasing in the two computed complexes (**h** and **g** cases). The reason is related to the weak H-bond interaction between glucose and cluster, but serves essentially to saturate the opening site in the cluster.

By comparing the optimized structures (**e**, **f**, **g** and **h**) to (**a**, **b**, **c** and **d**) clusters (before and upon glucose addition), we do not observe any substantial changes in terms of Sb-O bonds or Sb-O-Sb angles; instead the new created intramolecular H…O_1_ and H…O_2_ bonds between cysteine and the cluster (**SI**-**Table 1**). Again, the model structure (**e**) seems the most stable. Moreover the Gibbs free energies range from 8.81 to 31.77 kcal/mol, favoring the bridged metal-metal **e**. The solvation of (**e**, **f**, **g** and **h**) clusters are determined by using the SMD model (continuum solvation model). Solvation keeps almost same energy order, which varies slightly above **i** from 6.27 to 12.55 kcal/mol but contributes substantially to an enhancement of the global stability of these conformers (**SI**-**Table 1**). According to DFT-D3 results, Sb-Sb bonds were decreased from 3.153 Å to 3.021 Å (**SI**-**Table 1**, first column). Intermolecular connectivity remains the same because the glucose serve only to keep the cluster environment related to the small charges modified during the addition process.

The next modeling consist in the use of new hypothetical cluster (**i**, **j**, **k** and **l**) (**Fig. 2**), which are built upon cysteine addition on the last optimized clusters (**e, f, h** and **g**). Relevant optimized structures of (**i**, **j**, **k** and **l**) in gas phase calculated at B3LYP/LANL2DZ, are shown in **Table 2**. We observe then different intramolecular cluster-cysteine through H-bond links, varying from 1.491 Å - 1.631 Å mostly for O-H bonds in (**i** and **j**) and as well as 1.721 - 1.681 Å for N-H bonds (**Table 2**). These newly created O-H bonds show present standard H-bonds or Van der Waals type interactions between oxygens of the cluster or N-H. For all computed (**i**, **j**, **k** and **l**) tautomers, the Sb-Sb bond distances were increased from 3.143 Å to 3.551 Å. Regard to the binding energy computed in **i**, cysteine addition shows good inhibition (less than 18.6 kcal/mol). The binding energy can be evaluated by the Gibbs free energy related to the closest cysteine molecule and without any S-S bond breaking. For all optimized tautomers, the binding energies of glucose-antimony-cysteine are in the range of 18 - 24 kcal/mol through a new O-H and N-H bonds from the amine function group. In terms of dipeptide cluster interaction, the S-S bonds remain strong, closer to 2.283 Å without any dissociating or chemisorption of cysteine.

**Fig 2:**
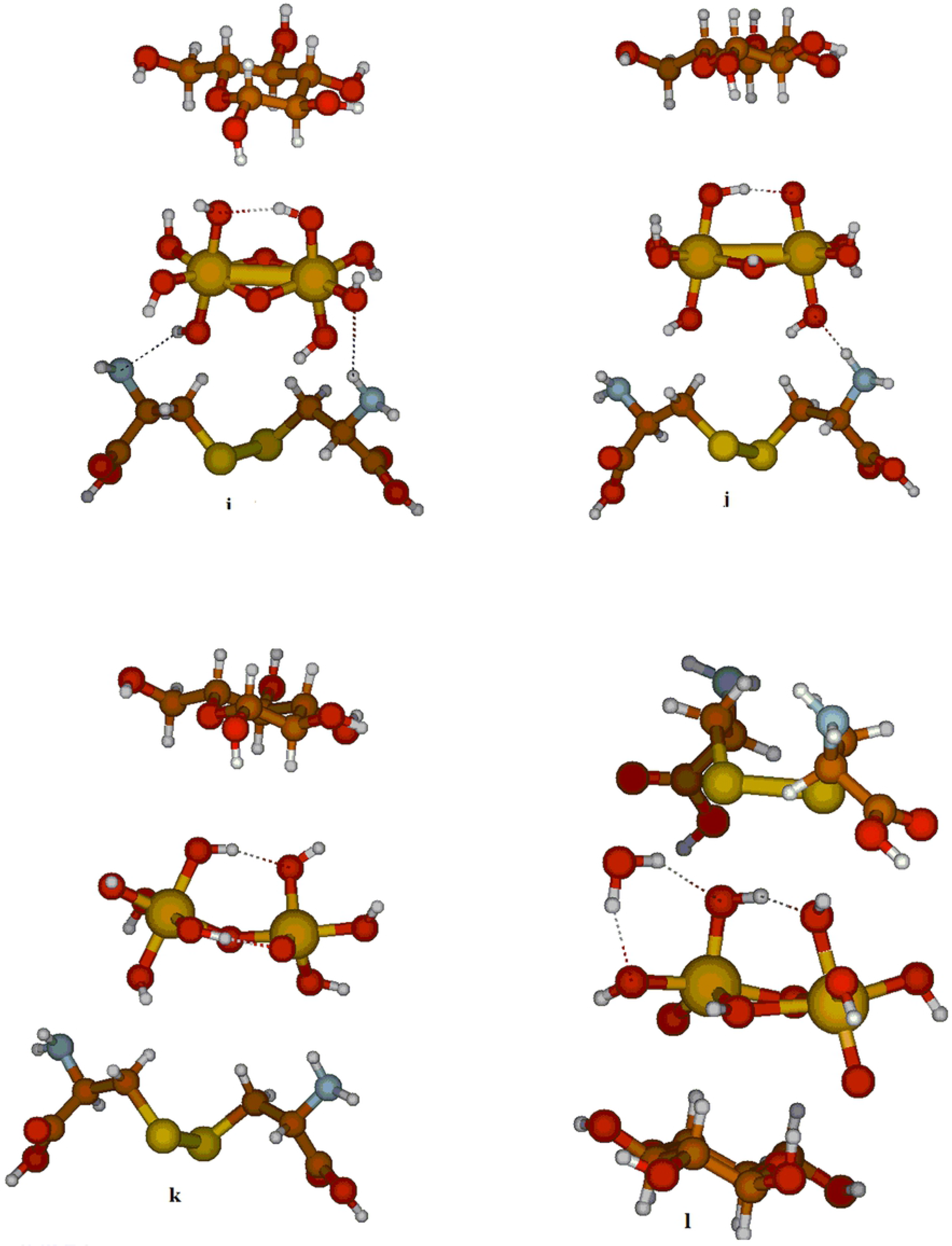
Optimized structures at B3LYP/Lanl2dz **(e, f, c** and **h** + glucose).

**Table 2.** Optimized structures of (**i**, **j**, **k** and **l**) calculated at B3LYP/Lanl2dz in gas phase and **f** calculated at **TPSS/Def2-TZVP (DFT-GD3BJ)**. (Bonds are in Angstrom and angles are in degrees). O_1_ and O_2_ are the closest oxygen pointing to glycose. O_3_, O_4_ are pointing from cysteine. ΔE: Total and Gibbs free energy changes ΔG (kcal/mol)/ in gas and water solution. Binding energy (BE), and ΔE are in (kcal/mol).

| **Tautomer +**<br>**Glucose +**<br>**Cysteine** | **i** | **j** | **k** | **l** | **(a) +**<br>**Glucose +**<br>**Cysteine**<br>**f**<br>**DFT- GD3BJ** |
| --- | --- | --- | --- | --- | --- |
| Sb-Sb | 3.145 | 3.261 | 3.609 | 3.551 | 3.027 |
| S-S | 2.283 | 2.274 | 2.283 | 2.283 | 2.053 |
| H----O<sub>1</sub> | 1.865 | 1.879 | 1.842 | 1.970 | 1.773 |
| H----O<sub>2</sub> | 1.622 | 1.878 | 1.711 | 1.589 | 1.915 |
| H----O<sub>3</sub> | 1.631 | 1.721 | 2.314 | 2.271 | 1.716 |
| H----O<sub>4</sub> | 1.493 | 1.581 | 2.617 | 2.104 | 1.852 |
| N---H | 1.72 | 1.682 | 2.100 | 1.998 | 1.991 |
| ΔE | 0. | 18.61 | 21.26 | 24.04 | 15.60 |
| ΔG Gas phase | 0. | 6.27 | 13.89 | 31.77 | 37.10 |
| ΔG (SMD=water) | 0. | 6.30 | 16.82 | 32.50 | 31.26 |
| (BE) without BSSE | -11.10 | -6.19 | -10.69 | -6.01 | -11.50 |
| (BE) with BSSE | -11.08 | -6.18 | -9.67 | -5.99 | -9.48 |

The total energy calculation of (**i**, **j**, **k** and **l**) with or without basis set super position error (BSSE) correction is summarized in **Table 2**, (last 2 rows). Effectively, this BSSE correction contributes to lower the energy of those optimized structures by a few kcal/mol, ranging between −11.08, −6.19, −9.67 and −5.99 kcal/mol respectively for (**i**, **j**, **k** and **l**) clusters.

Our calculations suggest a good interaction of cluster/dipeptide due to the intramolecular chelate ligands (amine or carboxyl group). Therefore, we propose antimony oxide can act as good inhibitor and block all sites without any dissociation of the molecules. The DFT-D3 correction with TPSS/Def2-TZVP, gives in all cases improved results in comparison to the B3LYP-LANL2DZ, particularly the shortness of S-S and C-S bonds. However, in contrast the B3LYP-LANL2DZ results, better agreement of cluster−dipeptide distance (intramolecular distances) with D3 correlation [36] (**Table 2, last column**). Furthermore, DFT-D3 correction improve the connectivity between atoms at long distance due to the acid-base interaction. And, it contributes to lower the energy of the optimized **f** structures by a 15.60 kcal/mol in term of total stability. Then le process of inhibition can occur easily.

For comparison of the computed most stable compounds in gas phase shown in **Table 1, 2, and SI-Table 1**, we summarized in **SI-Table 2**, the calculated charges, energy gap, and the dipolar moment of the most stable cluster (**a**, **e** and **i**) at different of level of theory. Therefore, the antimony charges remains positive in **i**showing their hard acidity during the full addition process whereas the oxygen atoms and nitrogen draining from the cysteine remains more negative (−0.63 e) vs (−0.75 at DFT-D3). Moreover, the large HOMO-LUMO gap preserves good stability of **i** with a lower dipole moment equal to 1.38 D vs 1.78 D computed at DFT-D3 This polarity is related to the cluster-ligand acid base reaction based on the proton mobility (OH from Sb oxide) and favors an attractive effect between cluster and cysteine (particularly with amino-acid groups). In addition, the cluster **i** is predicted to be the most stable compound, by 18 kcal/mol (**Table 3**). So far, the electrostatic forces between cluster surface and dipeptides show good interaction between nitrogen amino-acid less than the carboxylate groups. Interestingly, it seems in accordance with our calculation that the interaction donor (ligand) with the acceptor (cluster with empty <u>d orbitals</u>) plays an essential role in the inhibition process. When, the HOMO-LUMO energy levels difference between donor and acceptor is lower, this is opportune to realize a good inhibition for the studied system [40,41]. Also, the HOMO-LUMO wave function localization given at DFT-D3 contributes to make an overlapping through H-atom mobility leading a good interaction surface/substrate thorough (HOMO donor and LUMO acceptor and vice versa.

### Cluster modelling

Cluster models have drawn great attention due to their fundamental interest in basic research as well as the possibility of constructing nanostructured materials using such clusters as building blocks [37–41]. To simulate the inhibitor molecules effect using large cluster and large molecules in the antimony oxide interaction, calculations were performed on the big cluster/dipeptide. This theoretical approach has also been used in our previous studies of the interaction of organic molecules with oxides surfaces [40–43]. To achieve a better understanding of proposed mechanistic of the inhibition reaction process of cluster/dipeptides, the focus in this investigation is on the reactivity of trypanthione with the large neutral cluster [Sb_7_O_28_H_21_] **B** in both vacuum and water medium including glucose adducts.

### Molecular structures of [Sb_7_O_28_H_21_], trypanothione and glucose complexes

The neutral cluster structure [Sb_7_O_28_H_21_] **B,** was taken from Sb_2_O_5_ oxide structure experimental data [18] by adding hydrogen atoms to reproduce water environment. The molecular structure of [Sb_7_O_28_H_21_] **B**, was optimized at B3LYP/LANL2DZ level and reoptimized at DFT-D3. The Cartesian coordinates are given in the supplementary information and the molecular optimized structure are drawn in **Fig 3**. In **B**, the antimony center keeps an octahedral environment surrounded by different mono- or bi- or tri- bridging oxygens. The optimized parameters of **B** are listed in **SI-Table 3**. At B3LYP, the Sb-O bond distances are more elongated than at DFT-B3, and varying from 1.948 Å to and 2.052 Å versus 1.913 Å to 2.055 Å resepectively, whereas the O-Sb-O angle are ranging respectively at B3LYP for the terminal, bridging and tridendate oxygen from 117/121(°), 112/110(°), 110/137(°) vs at DFT-D3 108/106(°), 106/105(°), 128/109(°) repectively. Also between the two computed functional, the correlation implies a decreasing of bonds and angles and agreeing the X-Ray data [18]. With regard to the mutual interaction of Sb and O atoms in **B,** and their electrostatic affinity, we obtain antimony positive Mulliken charges 2.36 / 2.17 *e* and the negative oxygens −0.80 / −0.91 *e* and −1.07 / − 0.78 *e* for O_b_ and O_t_ respectively, and similar at DFT-D3 (see **SI-Table 3**). The O and Sb charges influence and enhance the polarizability of the terminal hydroxyl donor group (which will be considered as the cluster interface). Therefore, the cluster acidity (OH groups) plays also a crucial role in the reactivity cluster-dipeptide (see vide-infra). It is believed that the acidity constant (pka) of such metal oxides should be the key of the reactivity with dipeptide [44].

**Fig 3:**
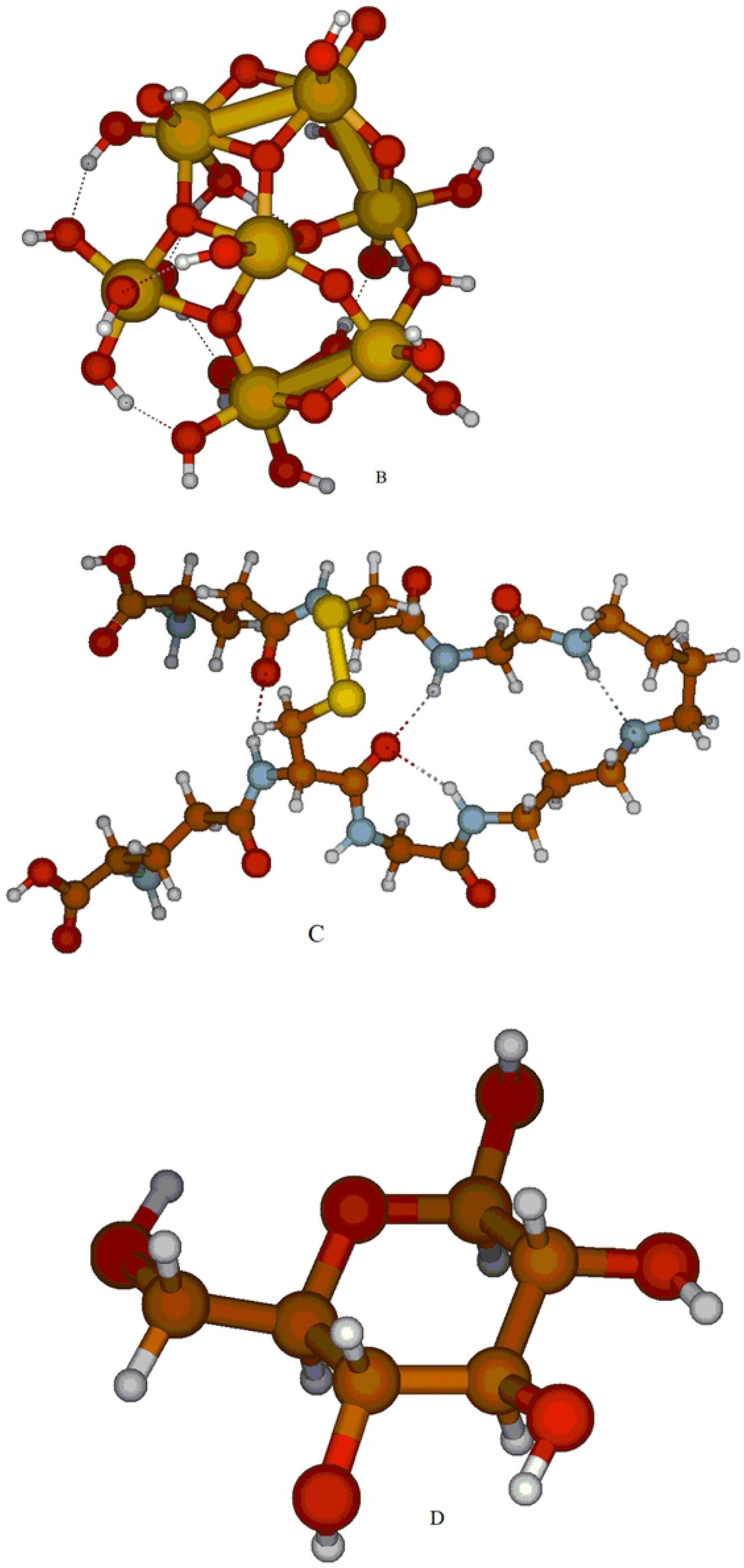
Optimized structures of (**i, j, k** and **l**) + glucose + cysteine, coordinated to the four optimized antimony tautomers at B3LYP/LANL2DZ.

We know also that the highest occupied molecular orbital (HOMO), the outermost orbital filled by electrons, is to be considered as an electron donor, while the lowest unoccupied molecular orbital (LUMO) is considered as an electron acceptor. A large HOMO-LUMO gap energy was seen for **B** in gas or water medium, limited between −8.41/−3.81 eV or −8.62/−4.07 eV respectively higher than calculated at DFT-D3. The high dipolar moment in solution is equal to 8.62/−4.07 versus −7.78/−4.56 D at DFT-D3, pointing to a good electrostatic interaction yielding an elongation of Sb-O bonds [45–46].

The trypanothione **C** molecular structure was also optimized at the same level of theory done for **B**. Structural Optimizations are drawn in **Fig 3**, middle. Metric parameters of **C** are presented in **SI-Table 3**. The S-S and N-H bond lengths are equal to 2.288 Å to 1.018 Å respectively versus 2.1051 Å and 1.241 Å, too short at DFT-D3. The C=O bonds are ranging from 1.263/1.271 Å. The S-C bond lengths are equal to 1.89/1.90 Å. In gas phase, the HOMO-LUMO gap of **C** is ranging between −5.72/−2.87 eV, as well as in aqueous medium, it is limited between −6.28 to −3.81 eV. The calculated dipole moment is 4.58 D. We conclude that those frontier orbitals (HOMO-LUMO) are in the same energy range of **B** and **C** and should react strongly [42] in term of overlapping. Then the inhibition will also dependent ultimately on these quantum parameters for both used methods.

The cyclic Glycose **D** molecular structure shown in **Fig 3**, right hand, is also considered as the most stable computed molecule regards to our calculation optimized at B3LYP/LANL2DZ and DFT-D3. Averages of C-O and C-C bond lengths in **D**, vary from 1.543 Å to 1.478 Å (see **SI-Table 3**, third column). The large HOMO-LUMO gap predicts a good stability of **C** ranging between −7.520/0.017 eV and −7.510/0.084 eV in gas phase and water environment respectively but higher in energy level at DFT-D3. But according to the difference established between HOMO-LUMO energy levels in **B** and **D**, one can say that the interaction became weaker than **C**.

### Optimised molecular structures of [Sb_7_O_28_H_21_]/glucose and [Sb_7_O_28_H_21_]/trypanothione and [Sb_7_O_28_H_21_]/glucose-trypanothione in gas phase and aqueous medium

To achieve a better understanding of proposed mechanism of inhibition by the cluster/dipeptide, we focus in our study on the reaction of trypanthione with [Sb_7_O_28_H_21_] giving **B1** drawn in **Fig. 4**, left hand. Most Sb-O distances are in the range 2.071/2.102 Å, while the O-Sb-O angle are ranging respectively for the terminal, bridging and tridendate oxygen from 111/108 (°) and 126-128 (°). The angles are more open, connected to the strong interaction between cluster **B** and trypanthione. The important revelation is the inter-molecular connectivity trypanthione/cluster via N-H or CO groups, and the easiest hydrogen mobility, considered as good chelating adducts. The distances we calculate are, respectively, equal to 1.613 Å and 1.682-1.78 Å, very short bonds. The S-S bond remains equal to 2.285 Å without any bond breaking or oxidative addition. Some of Sb-Sb bonds are increasing from to 3.362 Å to 3.581 Å, as well as some much longer Sb-O--H contacts. Mulliken charges for all atoms near the surface-surfactant interface decreased slightly (see **SI-Table 4**).

**Fig 4:**
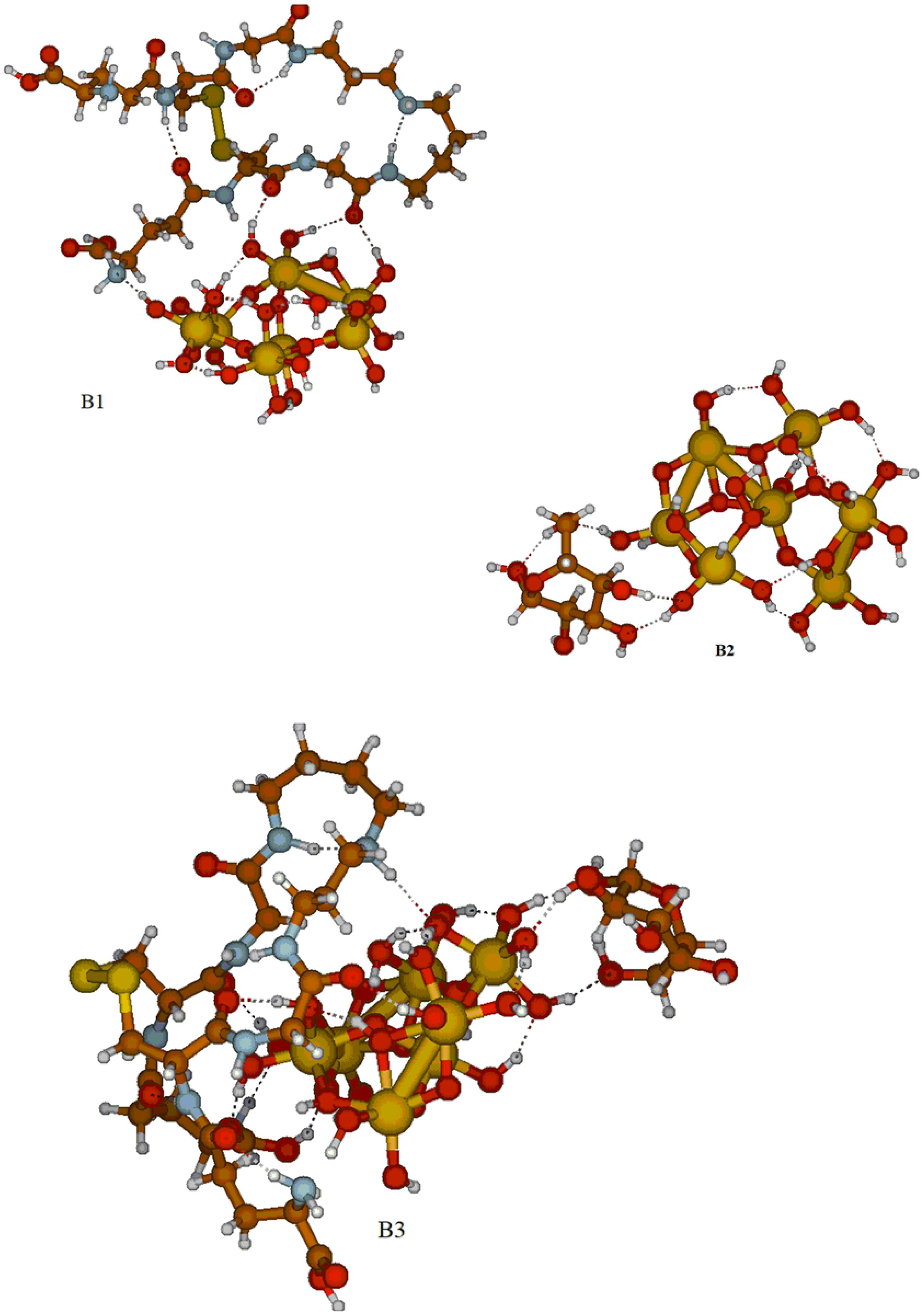
Molecular structure clusters optimized at B3LYP/ LANL2DZ level [Sb_7_O_28_H_21_]/trypanthione **B1,** [Sb_7_O_28_H_21_]/glucose **B2,** [Sb_7_O_28_H_21_]/glucose-trypanothione **B3,** The Sb atoms are shown in yellow spheres, and the O atoms in red, N atoms in blue, Carbon atoms in grey and the H atoms in white.

A large gap energy of HOMO-LUMO −6.39/−3.45 eV is observed in the gas phase versus −7.56/−4.13 eV in aqueous medium. This is due to the good interaction cluster/trypanothione, which shows a chelate molecule with amino-acid and carboxylate groups (i.e. (N-H-O) bond equal to 1.613 Å and (C-O-H) equal to 1.682-1.781 Å; very tight hydrogen bonds occur (see **SI-Table 4**). The stronger tendency towards deprotonation of the carboxylic group adsorbed in the symmetric bidentate mode can be explained by resonance: the symmetry between the two O atoms in the binding COO^−^ group leads to a stabilizing delocalization of the negative charge.

Following our computational results, the HOMO-LUMO energy difference is 3.10 eV, expecting a relative stability for **B1.** Although, the S-S bonds are strong, ΔG_**B1**_(Gibbs free energy) values are predicted to have good thermodynamic stability yielding 9.66/3.64 kcal/mol in gas phase and quantum solvation respectively. We observe that the trypanothione involving in inhibition process is not dissociated. The calculated dipole moment calculated for **B1** are large, 14.16 and 14.46 D in gas phase and solvated medium respectively.

We calculated also the binding energies of the glucose in **B2**, which is lower by 2.50/1.50 kcal/mol than **B1** (**SI-Table 4, Fig 4**) revealing an endothermic and spontaneous chemical process. We observe also a large gap HOMO-LUMO ranging from −7.35 to −4.33 in vacuum and −7.56/−4.15 in aqueous medium. Therefore, smaller hydrogen contact was seen in B2 than in B1 related to the energetic gap between HOMO-LUMO fragments. The partial intermolecular bonding interaction depends on the linking hydrogen bonding (H-bonds). This type of interaction is seen in such compounds. It involves a hydrogen atom located between a pair of other atoms having a high affinity for electrons; such a bond is weaker than an ionic or covalent bond but stronger than van der Waals forces. Hydrogen bonds can exist between atoms in different molecules or in parts of the same molecule, as noted here with weak intermolecular hydrogen bond.

To get insight into the dipeptide **C**/cluster **B** interaction in vacuum or in solution, we consider a new large cluster model **B3** coordinated simultaneously to the glucose and the trypanothione molecule. Calculations were performed in both three fragments, **C**, **B**, and **D**, by examining the optimized geometry computed at B3LYP/LANL2DZ level. The molecular structure of **B3** is drawn in **Fig 4**. The calculation shows an elongated Sb-O distances varying from 1.993 Å and 2.151 Å whereas the O-Sb-O angle are ranging from 105-142°.

The next revelation is that we observe intermolecular bonding through short H-bond (amine) and the ketonic bonds. These two short non-bonding distances are calculated respectively equal to 1.591/1.671 Å, (**SI-Table 4**, third column). Mulliken charges remain without large changes. The HOMO-LUMO gap is between −6.66/−1.48 eV in gas phase and −6.30/−3.91 eV in water solution, suggesting a good stability of the bulk. The binding energies of the cluster to the solvated cluster surfactant molecules were calculated at the same level of theory, equal to 18.45 and 3.1 kcal/mol respectively in gas and water medium. It consists a tight intramolecular interaction, favored by a short N-H distance from amino-acid and good trypanothione chelating contributing to metal inhibition. This option is related to the closer HOMO-LOMO energy level of interacting cluster **B** and trypanothione. Moreover, the amine and carboxylate groups closer to the oxide surface (Sb_2_O_5_,nH_2_O). Parallel attack of trypanothione wil sulfur lone pairs close to the surface is predicted to be by less than 6 kcal/mol [43]. The cluster trypanothione frontier orbital (FO) is localized mainly in the nitrogen and carbonyl groups (CO). This interface binding site of Sb atoms to TR **C** with bound Sb represents a good hydrophilic metal-ligand interaction in solution. The formation of reaction cluster **B**/dipeptide **C** is quasi- reversible with lower ΔG. Therefore, we suggest that the oxide (Sb_2_O_5_.nH_2_O) is a potent inhibitor. According to the parameters calculated in this study, Antimony oxide should enhance the inhibition system, by leading a good stability of cluster/dipeptide. The nitrogen charges are slightly negative of −0.65/−0.41 e, depending of the coordination site, whereas the total charge of the antimony cluster remains the same. The Sb-O-H-Ligand bond seems to be governed by electrostatic attractions leading to large Sb^V^ ← σ_N-H_ donation. These calculations provide information about the strength of the Sb-O-H bond and the interactions between Sb and hydroxyl group as a function of the σ or π donor/acceptor process. The cluster formation energy per Sb-O unit and the electron affinity tend to increase, whereas the ionization potential tends to decrease with the cluster size *n*.

By using the SMD continuum modeling of the water environment [40], some structural modifications occur, increasing the length of certain Sb-O bonds, an increasing of other Sb-O bonds lengths (see second columns of **SI-Table 4**). We observe a large polarization of the cluster indicating the antimonic acid (Sb_2_O_5_·nH_2_O) exhibits a high protic conductivity [89]. According to the lowest computed ΔG, we predict a stable the complex with a good inhibition efficiency. The dipole moment remains also higher in term of polarity, and favor good interaction surface/substrate. The stabilization of the solute–solvent interaction and the lowering of non-electrostatic energy for the protonated complexes indicate that the adsorption might involve chemisorption, which is energetically stabilized by donor– acceptor interactions between lone pair electrons of the surfactant molecule and the vacant d orbitals of the Sb atoms. The SMD [44] calculations show that the relative energies of the computed complexes, are lower when increasing the polarity of the solvent and size of the molecule. The solvation of the surfactant molecules provides significant changes in the charge distribution, giving rise to a larger dipole moment than that obtained in the gas-phase calculation. This is event is enough to make easy the protonation of the surfactant. Then, it seems the acidity and basicity of the cluster play a big role, depending of the number of H_2_O molecules surrounding the cluster and the acidity constant (pka) which can easily modify the cluster-dipeptide reactivity [47], whereas a more general treatment should be based on electron donor-acceptor (Lewis acid-base interactions).

In this section, the use of modern dispersion-corrected DFT is able to correctly describe various types of van der Waals interactions. Structural improvement are seen in Sb-O bond lengths, and Sb-O-H-N interaction particularly. The D3 correction gives in all cases improved results in comparison to B3LYP (see **SI-Table 4**), particularly in Sb-Sb bonds and intramolecular distances (i.e. Sb-O---H---N and Sb-O---H---O). By using the dispersion correction more stability of the full complexes (cluster/dipeptide) was observed by (10-20 Kcal/mol).

The addition of the dispersion correction to the DFT results improves the outcome significantly. S-S bonds and bond energy were decreased contrary to the B3LYP results. Large umbrella of H-bonds connection in cluster/dipeptide were added with a distances ranging from 1.639 to 1.871 Å adding more chelated effect. This effect is emphasizing in the easiest chelating molecule surrounding the cluster by creating the large hydrogen interaction and contributing to decreases the intramolecular interactions by making a short Sb-O-H----N or Sb-O----H---O, type van der Waals bonds. This kind of interaction are the keys of the inhibition depending of H-bonds formed during the process the complexation (See in the Supporting Information, **SI-Table 4**. However, in contrast the DFT-D3 results are in better agreement of cluster−dipeptide stability (intramolecular distances) compared to B3LYP correlation [36] with a good binding energy 30 kcal/mol vs 18 Kcal/mol computed with B3LYP) (**SI-Table 4**).

Dispersion-corrected DFT-D3 gives a wide variety of hydrogen interactions in extra in such model systems in the vacuum and in solution and the complex inter- and intramolecular interactions present in the bulk (clustering model). This lead to an arising inhibition process.

### Molecular electrostatic potential (MEP)

The molecular electrostatic potential (MEP) profile is a crucial tool to grasp the molecular interactions in a given molecule. It is very useful for interpreting and predicting relative reactivity sites for electrophilic and nucleophilic attack, hydrogen bonding interactions studies of zeolite, molecular cluster and crystal behavior investigation of biological recognition, and the correlation and prediction of a wide range of macroscopic properties.

The 3D plots of the MEP of **B**, **C** and **B3** [Sb_7_O_28_H_21_]/glucose-trypanothione are shown in **Fig. 4**, which was the optimized molecular structure at the B3LYP/6-311G++(d,p) level basis set. Fig 5 shows the electrostatic potentials at the surface for clusters, using different colors with most important areas of interaction and hydrogen interactions. It illustrates additional insights on the associated nature of attractive forces based on molecular electrostatic potential (MEP) computations. The red color parts represent the regions of negative electrostatic potential, the blue lines represents the regions of positive electrostatic potential and the green color parts show the area of low values potential. Negative electrostatic potential regions are those that are likely to attract protons, while positive electrostatic potential corresponds to regions of repulsion of the proton by the atomic nuclei in regions where electron density low exists and the nuclear charge is incompletely shielded. The potential increases in the order < orange < yellow < green < blue. The analysis of the MEP of **B3** exhibits very good agreement with the calculated data illustrating additional insights on the associated nature of attractive forces based on molecular electrostatic potential (MEP) computations. The MEP maps reveal the around O and N atoms negative (red and yellow) regions for both compounds, and around the nitrogen atom from **C** and especially the around the N –atom in trypanothione as the positive (red) region and support the presence of hydrogen bonds.

**Fig 5:**
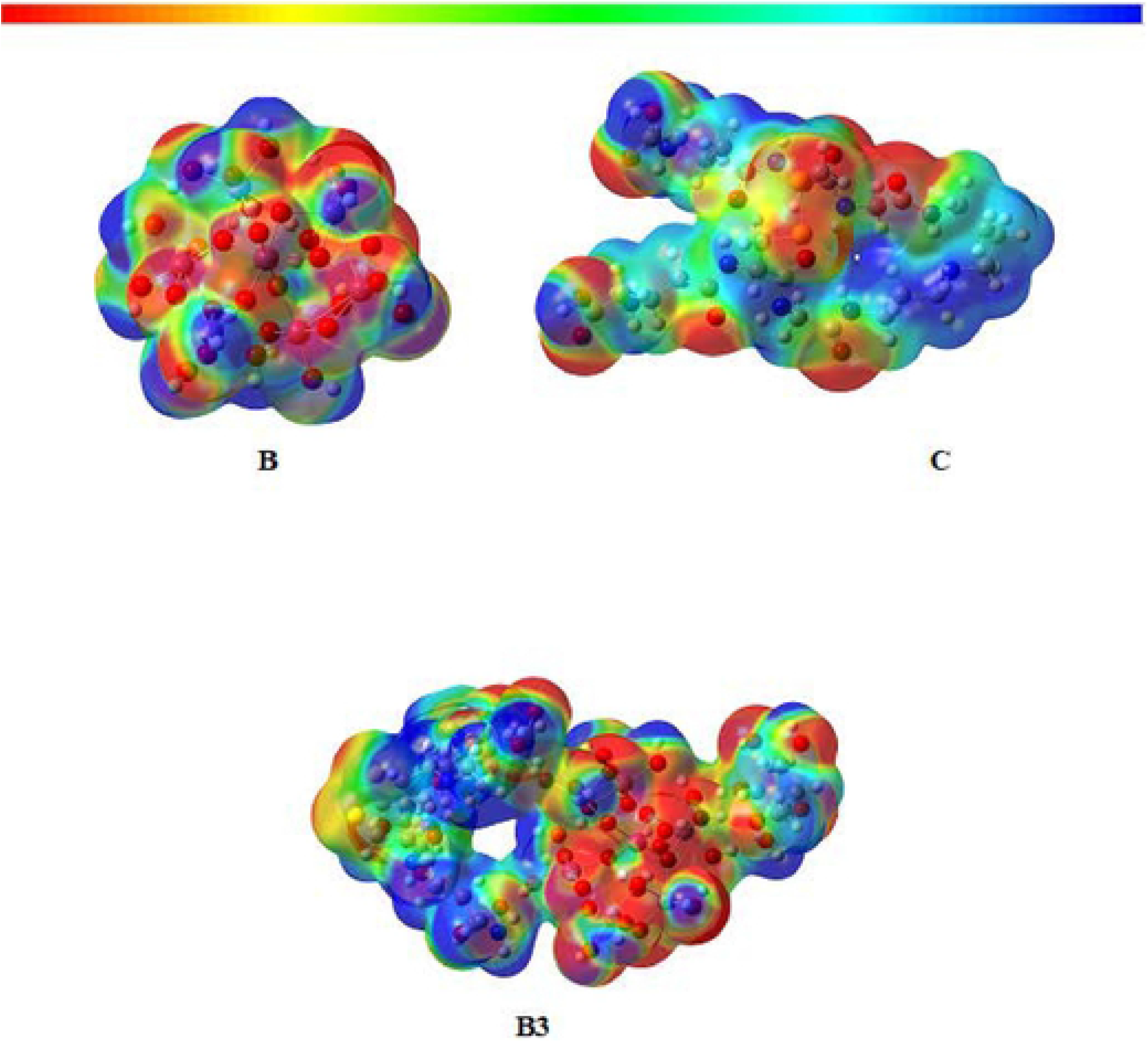
Illustration The molecular electrostatic potential and negative region in space (ESP) computed - at B3LYP/6-311G++(d,p) of [Sb_7_O_28_H_21_]/ B, C trypanothione. and **B3** [Sb_7_O_28_H_21_]/glucose-trypanothione. MEP surfaces are plotted in two manners, either mapped on electron density surfaces (rainbow plots with the color scale shown; isodensity = 0.0004) or positive (blue) and negative (red) regions in space (range = ±2.2 au; isodensity = 0.02). Atomic color code in the molecular structures: Sb, red-, larger spheres; C, gray; N, blue; O, pink-light small spheres; H, light-blue, smaller spheres and S, yellow.

## Conclusions

Relative stabilities were explored in different clusters coordinated (or not) to the cysteine or trypanothione of the one layer or two layer models of (Sb_2_O_5_.nH_2_O), at a reasonably high level of DFT and DFT-D3 computations. Calculations show the energy stability of different conformations, corresponding to dipeptide/cluster attraction is computed by a few kcal/mol. The reason is related to the good interaction between HOMO and LUMO of the fragment metal oxide/dipeptide and the good donor-acceptor character of the system. The theoretical analysis here reported on reduced the trypanothione reductase (TR) in a complex with (Sb_2_O_5_.nH_2_O) shows a cluster is good candidate for TR inhibiting. The structural and thermodynamic parameters analysis clearly demonstrates that inhibition should occur with a good acidic cluster, with large of hydroxyl groups. According to our Mulliken charges, the electrostatic forces between cluster atoms and inhibitors sites are strong related to the acidity of Sb_2_O_5_ with contributions from creation of dipole-dipole interaction through Hydrogen bonding. In the respect of the dipeptide reaction with cluster, NH_2_ can take up easily several proton from the oxide surface via proton mobility, modulated by an enhancement of electrostatic effects with a good sigma ligand donor. According to the computed dipole moment, high polarizable character of the cluster was observed in solution. The authors strongly believe that the use of a novel approach, based on protein-encapsulated metals, may represent a promising strategy for the treatment of cutaneous Leishmaniasis. To test the antiparasitic activity on Leishmania cells, we will used AgNPs or AuNPs as drug-delivery agents (calculations are underway) to continue this pertinent study.

## Acknowledgement

H.R. acknowledges the European community for supporting this work through the project H2020, and special thanks to the Arab Fund from Kuwait. Helpful discussion and suggestions were obtained from Prof. T. Cundry, Prof. M. Omary (UNT) and Prof R. Hoffmann (Cornell University). The computing facilities Finnish IT Center for Science (CSC) and American CASCaM, UNT, are acknowledged for computer time. Some feedbacks were done by Phd students, Zhou Lu, and Kurt Bodenstedt.

## Supporting Information Available

The optimized Cartesian coordinates at the B3LYP/LANL2DZ and TPSS/Def2-TZVP (DFT-GD3BJ) levels of theory, are given as Supporting Information.

## Supplementary information

**SI- Fig 1:**
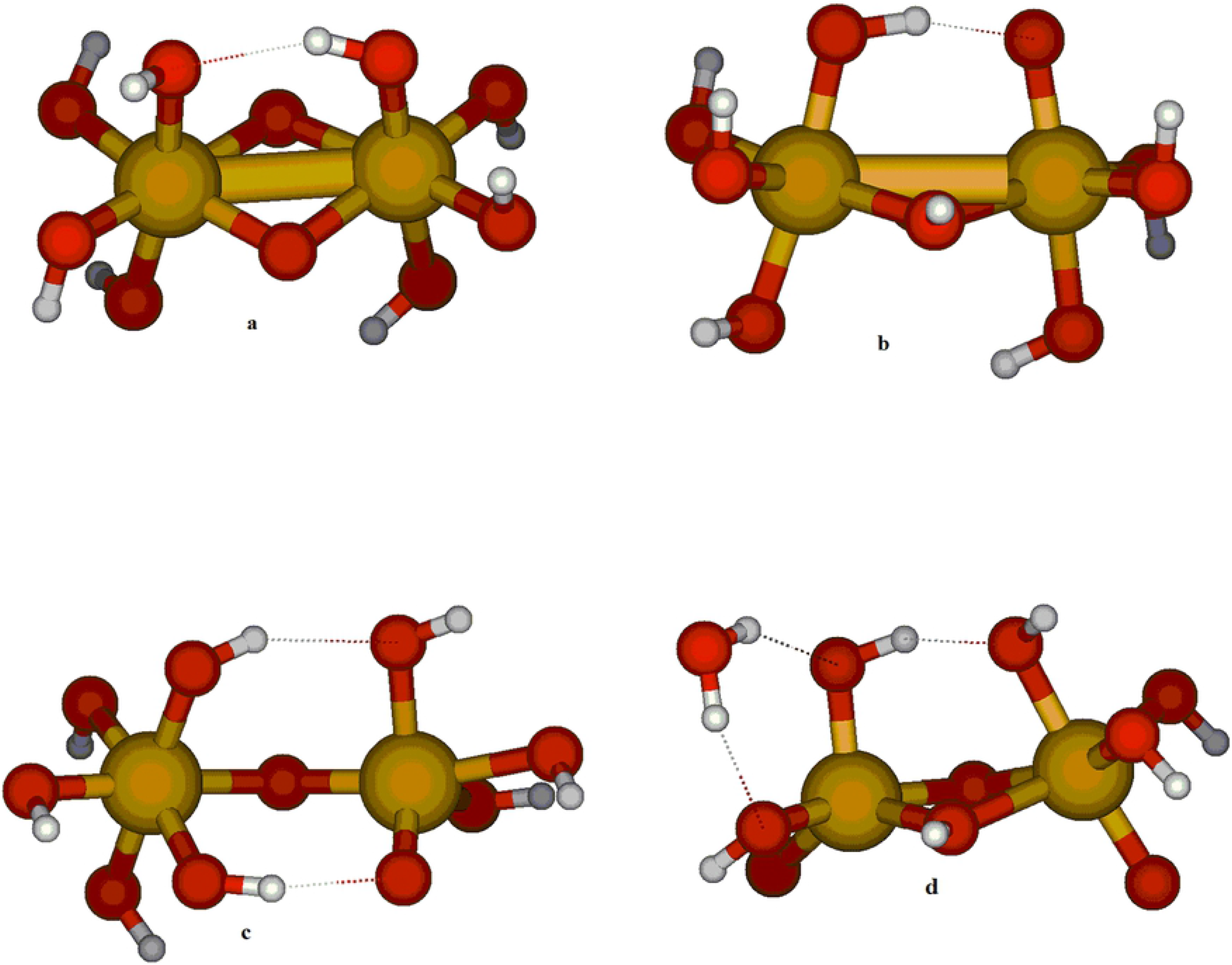
Molecular structures of different optimized tautomers of [(Sb_2_O_10_H_8_)]^−2^, **(a, b, c** and **d**). Sb: purple ball; O: red ball and gray ball: H.

**SI- Fig 2:**
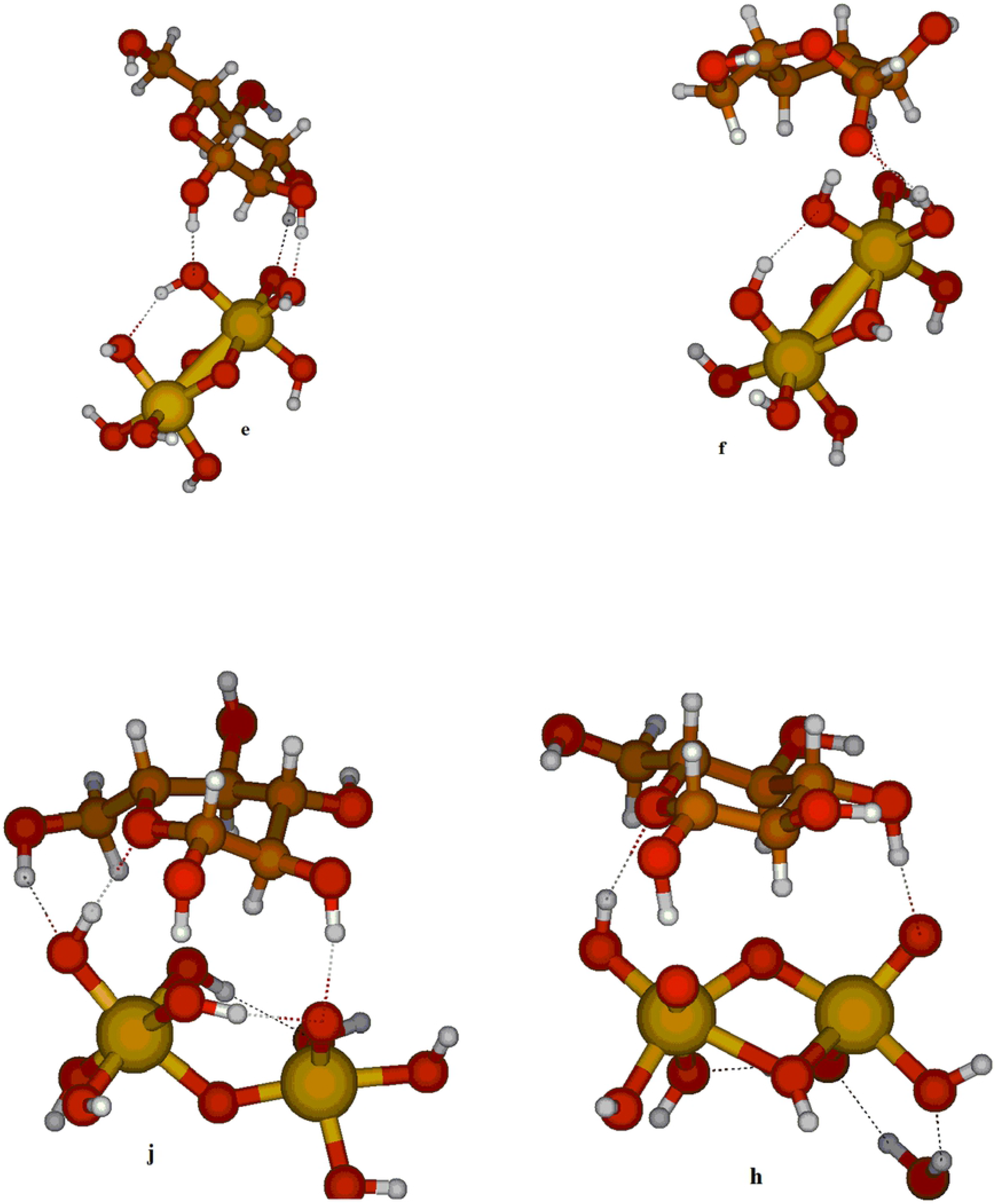
Molecular structure of **B** cluster of [Sb_7_O_28_H_21_], trypanthione **C** [C_27_H_47_N_9_O_10_S_2_], and glucose **D** [C_6_H_12_O_6_), optimized at B3LYP/ LANL2DZ level. The Sb atoms are shown in yellow spheres, and the O atoms in red, N atoms in blue, C atoms in grey and the H atoms in white.

